# DGAT-onco: A powerful method to detect oncogenes by integrating differential mutational analysis and functional impacts of somatic mutations

**DOI:** 10.1101/2020.02.15.947085

**Authors:** Haoyang Zhang, Junkang Wei, Zifeng Liu, Xun Liu, Yutian Chong, Yutong Lu, Huiying Zhao, Yuedong Yang

## Abstract

**Motivation:** Oncogenes are genes whose malfunctions play critical roles in cancer development, and their discovery is a major aim of cancer mechanisms study. By counting the mutation frequency, oncogenes have been identified with frequent mutations, while it is believed that many more oncogenes could be discovered by differential mutational profile analysis. However, it is common that current methods only utilize mutations in the cancer population, which have an obvious bias in background mutation modelling.

**Methods:** To predict oncogenes efficiently, we developed a method, DGAT-onco that analyzed the frequency distribution and functional impacts of mutations in both cancer and natural population. Our method can capture the mutational difference of two population, and provide a comprehensive view of genomics basis underlying cancer development. DGAT-onco was constructed by germline mutations from the 1000 Genomes project and somatic mutations of 33 cancer types from the Cancer Genome Atlas (TCGA) dataset. Its reliability was verified on an independent test set including 19 cancers from other sources.

**Results:** We demonstrated that our method is more effective than alternative methods in oncogenes discovering. Using this approach achieves higher classification performance in oncogene discovery than 6 alternative methods, and 22.8% significant genes identified by our method were verified as oncogenes by the Cancer Gene Census (CGC).

**Availability:** DGAT-onco is available at https://github.com/zhanghaoyang0/DGAT-onco.

## 1 Introduction

Oncogenes are genes that provide the growth advantage of a cell when activated by mutations. These genes have potential in cancer development, and their discovery may provide new insights in cancer diagnosis and therapies (Croce, 2008). Since functions of genes are associated with the impacts of mutations, it is believed that oncogenes can be recognized by detecting its specific mutation profile (Vogelstein, et al., 2013). However, apart from the most frequently mutated genes, it is difficult to determine new oncogenes because mutations are highly heterogeneous among individuals and different cancer types (Pon and Marra, 2015). Therefore, it is necessary to develop computational methods for the discovery.

High-throughput cancer genome sequencing consortia like the Cancer Genome Atlas (TCGA) (Zhu, et al., 2014) and International Cancer Genome Consortium (ICGC) (Chin, et al., 2011) provide comprehensive insights of oncogenomics, based on which numerous methods for oncogenes identification have been developed. For example, as an early attempt, MutSig (Forrest and Cavet, 2007) performs mutational significance analysis to identify oncogenes. This method assumes a uniform background mutation rate across all genomic positions. To measure the background mutation rate, it splits mutations into serval categories by their similarity and accumulates the probabilities of observed mutations in each sample. Thus, oncogenes can be identified with greater observed mutation probabilities than the background ones. Implementation of MutSig in the breast and colorectal cancers achieved higher accuracies than other methods without utilizing the mutation probability of total genes. Considering that the background mutation rates might change across all genomic regions, an improved method, MutSigCV (Lawrence, et al., 2013) estimates the background mutation rates based on silent mutations in each gene and non-coding mutations in its surrounding regions before frequency comparison. Additionally, it integrates covariates in DNA replication, transcriptional activity, and mutation frequency variance across patients in the analysis. As a result, MutSigCV discovered more oncogenes than its previous version.

Although these frequency-based methods successfully reveal cancer-related genes with a high frequency of mutations, such as TP53 and KRAS. Unfortunately, there are still two challenges ahead (Vogelstein, et al., 2013). First, it is difficult to identify genes with relatively few numbers of mutations but playing important roles in cancer development, because the background mutation varied among different patient and genomics regions. Second, since many mutations occurring in genes may not damage its functions, it is a challenge to detect and exclude them before analysis to avoid a bias. In order to solve the issue, it is necessary to consider the impacts of mutations for an accurate prediction. Thus, considering the clustering tendency of mutation in specific protein regions, Oncodrive-CLUST (Tamborero, et al., 2013) uses clustering scores to measure the clustering tendency in particular protein regions and synonymous mutations. In this method, genes with significant clustering bias with respect to synonymous mutations are regarded as oncogenes. Since Oncodriver-CLUST detects oncogenes with a different thought, its result had been proved to be a useful supplement of frequency-based methods. At the same time, with the development of pathogenic scoring approaches which prioritize the relationship between mutations and diseases, OncodriveFM (Gonzalez-Perez and Lopez-Bigas, 2012) has been developed to perform differential analysis based on gene functional impacts. The method uses SIFT (Ng and Henikoff, 2003), PolyPhen2 (Adzhubei, et al., 2010), and MutationAssessor (Reva, et al., 2011) to classify non-synonymous mutations into several functional impact groups and constructs a matrix to represent the functional impacts of a gene in all samples. Thus, the method could detect genes with a greater bias of functional impacts. Its applications to glioblastoma cancer, ovarian cancer and leukemia detected most of genes found by previous method and revealed several new pathways with high functional bias. Later, OncodriveFML (Mularoni, et al., 2016) had been developed by further improving the measurement of function impacts. This method splits genomic positions into different element and calculates their function impact bias with different formulas. Recently, a new method WITER(Jiang, et al., 2019) described a regression-based method in oncogenes discovery. Unlike the methods using existing mutation scoring systems, it pre-trained a score for each mutation based on Catalogue of Somatic Mutations in Cancer (COSMIC) data. Then, it used a stepwise procedure to derive the potential oncogenes one by one from a negative binomial model of mutations, until all genes in the model were non-significant. Moreover, in the datasets with a small sample size, it can also transfer the mutations of non-significant genes as background from large dataset to increase its statistical power.

Although these methods have been proved to be effective in oncogene detection, the background mutation they used were estimated from only cancer samples without consideration of actual distributions from the healthy. To address this issue, DiffMut utilizes germline mutations from natural populations as background (Przytycki and Singh, 2017) to boost the power. For each gene, DiffMut compares the distributions of mutation number between TCGA population and the 1000 Genomes population. The distribution difference is estimated by the unidirectional Earth Mover’s Difference (uEMD) (Rubner, et al., 2000), a statistics to measure the difference between two distributions or shapes. Because of the better estimation of the background mutation, DiffMut achieved higher discrimination than other methods. However, the method treated all mutations equally and neglected functional differences between mutations, which may lose power in mutation profile modelling.

In this study, we have developed a new method DGAT-onco by weighing mutations according to functional impacts and using a background from the germline mutations in the 1000 genome project. This was also inspired by our previous studies to develop DGAT series of tools for disclosing disease-related genes and pathways for genome-wide association studies study (Zhao, et al., 2016; Zhao, et al., 2017). By tested on 33 cancers from TCGA and 19 cancers form other sources, DGAT-onco shows better performance than other methods when validated on the Cancer Gene Census (CGC) known oncogenes list.

## 2 Methods

### 2.1. Mutation Data Collection and Annotation

TCGA dataset(Zhu, et al., 2014): all level 3 somatic mutation data was downloaded in September 2018 through four standardized variant calling pipelines (Muse, Mutect2, SomaticSniper and Varscan), each with 33 Mutation Annotation Format (MAF) files for 33 cancer types. After keeping mutations appearing in all of these four pipelines, we obtained a total of 2,024,409 mutations, namely TCGA dataset.

TS19 dataset: for the independent test, we downloaded somatic mutations of cancers from the cBioPortal (Cerami, et al., 2012) in November 2019. The cancer data was selected by the following criteria: 1) This cancer type is included in the TCGA dataset; 2) The sample size is the largest of its cancer type; and 3) Its source is not from the TCGA. Finally, we obtained a total of 445,249 mutations from 19 cancer types (Armenia, et al., 2018; Bailey, et al., 2016; Dulak, et al., 2013; Imielinski, et al., 2012; Johansson, et al., 2016; Johnson, et al., 2014; Jones, et al., 2014; Kim, et al., 2015; Krauthammer, et al., 2012; Lowery, et al., 2018; Pereira, et al., 2016; Reddy, et al., 2017; Sato, et al., 2013; Schulze, et al., 2015; Soumerai, et al., 2018; Stransky, et al., 2011; Tyner, et al., 2018; Vasaikar, et al., 2019; Wang, et al., 2014), namely TS19 dataset.

The 1000G dataset: the germline mutations were downloaded from the phase 3 whole-genome mutation data of the 1000 Genomes Project (released at 20130502) (Nature, 2015). The data consisted of 25 Variant Call Format (VCF) files for the 24 chromosomes and mitochondrial chromosome. We excluded variants on the Y chromosome as it only contained information of males, and the mitochondrial chromosome as the TCGA dataset did not have such information. Finally, we obtained 84, 739, 838 human genetic variants from 2504 individuals, namely the 1000G dataset.

All the mutations in the three datasets were annotated by the ANNOVAR (Wang, et al., 2010) to obtain pathogenicity scores according to the dbnsfp33a database (Liu, et al., 2016). For the TCGA and TS19 datasets, we used the provided annotation of gene names and mutation types in the analysis. For the 1000G dataset without annotations, the gene names and mutation types were annotated based on the refGene database (hg19 version) (O’Leary, et al., 2015). In the analyses, we kept only non-synonymous mutations that cause changes of coded amino acids. Table S1 detailed the 3 datasets.

### 2.2. Method Overview

DGAT-onco is a method to detect oncogenes with a significantly different mutational profile between somatic mutations in the cancer population and germline mutations in the natural population. In order to integrate the functional impacts of mutations, DGAT-onco assesses the mutational profile of each gene by summing the predicted pathogenic scores of all its contained mutations. Here, we used 23 types of pathogenic scores deposited in the dbnsfp33a database (Liu, et al., 2016), as detailed in Table S2. For comparison, we also used a constant scoring function to mimic the DiffMut, i.e., a gene profile as the number of mutations. Note that all these scores have been normalized to be in a range from 0 to 1, with a greater score indicating a higher probability to be damaging.

As shown in Fig. 1, for a population, the summing scores of genes were presented in a matrix *S* = [*s*_*ij*_], where *s*_*ij*_ is the summed functional impact scores over all mutations on gene *i* of individual *j, s*_*ij*_ can be computed as 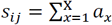, where X is the total number of non-synonymous mutations in gene *i* of individual *j*, and:

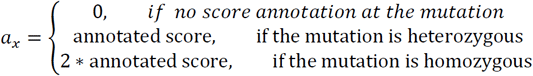

**Fig. 1.**
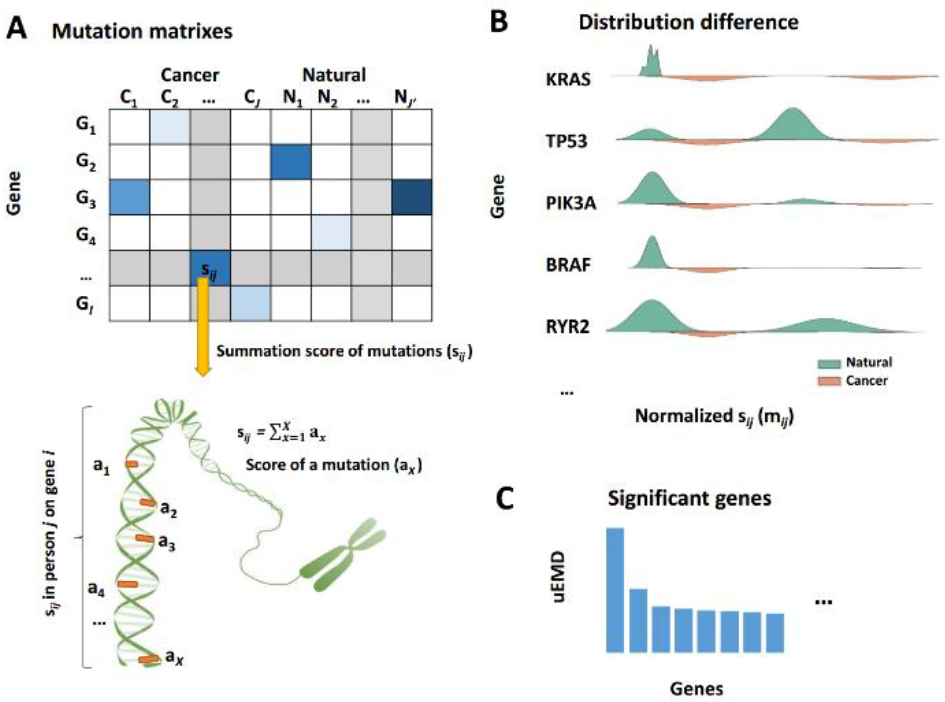
Overview of DGAT-onco framework. DGAT-onco evaluates the oncogenicity of each gene by comparing its mutational profile between cancer and natural populations. (A) The input of DGAT-onco was a mutation matrix under a specific pathogenic scoring function. In the matrix, we measured the summation of mutation score (*S*_*ij*_) of each person and each gene, before conducting a rank-based normalization. (B) For each gene, we extracted and compared its distributions of normalized *s*_*ij*_ (*m*_*ij*_) in natural and cancer population. (C) We measured the statistical difference between two distributions by uEMD and tested the significance.

Since TCGA dataset has not information about heterozygosity, its mutations are regarded as heterozygous. For 1000G dataset, all mutations in the X chromosome of males were considered to be homozygous. The matrix *S* was then normalized to a new matrix *M* with each value ranging from 0 to 1. Briefly, for each person *j*, all genes were ranked according to {*s*_*ij*_, *i*=1, ngene}, and *m*_*ij*_ for gene *i* was computed as its rank divided by the total number of genes. Based on the matrix *M* = [*m*_*ij*_], we evaluated the statistical distance of each gene between two populations by the uEMD, the same statistics method as previously used in DiffMut. In details, for a gene *i*, we divided the range 0 to 1 into 100 equal bins, and counted the proportions of {*m*_*ij*,_ *j*=1, npersons} falling in all bins. According to the proportions in all bins, the uEMD for each gene was computed as:

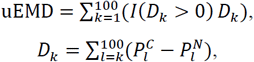

where 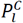and 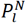are the proportions at the l bin for the cancer and natural individuals, respectively, and I() was an indicator function with output of 1 when the condition is true, otherwise 0.

To perform the statistical significant test for the obtained uEMD values, we calculated 5 random uEMD values by radomly shuffling the mutation ranks in each individual, and the P-value was computed by the one-sided one-sample t-tests. Q-value can be computed by the Benjamini and Hochberg method (Benjamini and Hochberg, 1995) as *P*\**n_test*/*rank*, where *rank* was the rank of a gene in the totally *n_test* genes according to P-values. All genes with Q-value < 0.05 were considered to be statistically significant. In addition, the largest P-value among these significant genes is taken as a threshold, and any gene with P-value less than the threshold is also considered significant, even if its Q-value is greater than 0.05 (Benjamini and Hochberg, 1995).

### 2.3. Methods to Compare

There have been multiple popular methods for oncogene detection. According to their evaluation studies, we selected 3 recent methods (WITER, ITER (Jiang, et al., 2019), and DiffMut (Przytycki and Singh, 2017) and 3 classic methods (OncodriveCLUST (Tamborero, et al., 2013), OncodriveFML (Mularoni, et al., 2016), and SomInaClust (Van den Eynden, et al., 2015)). Although DGAT-onco only need non-synonymous mutations, we provided all types of mutations for fair comparison.

The performance of a method was evaluated by the area under the precision-recall curve (AUPRC) and the precision of top genes detected by the method. A gene was regarded as an oncogene if it has been marked as an oncogene according to the somatic mutations in the Cancer Gene Census (CGC) database (https://cancer.sanger.ac.uk/census). Table S2 detailed the responding relationship of cancer names in the TCGA datasets and the CGC database, with a portion of names matched by reviewing literatures. The details of the usage of alternative methods and the measurement of AUPRC are described in Supplementary1 Notes.

### 2.4. Assessing the Enrichments of Genes on Functional Categories

The functions of genes identified by DGAT-onco were assessed by comparing these genes with numerous functional gene categories. These gene categories are defined by their role in biological processes, pathway membership, enzymatic function, and so on. The enrichment analysis was performed by Metascape (http://metascape.org/gp/index.html#/main/step1), an integrated portal leveraging knowledge from over 40 databases (Zhou, et al., 2019).

## 3 Result

### 3.1 Performances of DGAT-onco combined with Pathogenic scores

Based on each pathogenic scoring function for mutations, we generated a mutation profile for each gene in a person, through which mutational differential analysis was conducted to calculate the statistical difference (uEMD) of the gene between the TCGA and natural populations. The genes with high uEMD were considered to be candidate oncogenes. Fig. 2A shows the average AUPRC over 33 cancer types for DGAT-onco combined with 24 scoring functions (23 pathogenic scores and one constant score). The highest AUPRC of 0.208 was achieved by the one with the Mendelian Clinically Applicable Pathogenicity (M-CAP) (Jagadeesh, et al., 2016), a score function trained from multiple pathogenicity scores by a gradient boosting tree model. The model with MetaLR ranked the 2^nd^ with an AUPRC of 0.192 that was 8% lower than the one with M-CAP. The model with phastCons20way ranked the 3^rd^ with an AUPRC of 0.180, which is significantly lower than the one with M-CAP according to the paired t-test (*P*<0.01). Other 21 scoring functions resulted in AUPRC values ranging from 0.13 to 0.178, all significantly lower than the one with M-CAP by the paired t-tests. The one with the constant scoring function, which has been designed to mimic DiffMut method, ranked the 7^th^ among the 24 scoring functions. The corresponding AUPRC was 0.175 that is 17% lower than the one by DGAT-onco with M-CAP function. Fig. S1 details the AUPRC values of 24 scoring functions in a heatmap. Direct comparisons in Fig. 2B shows that the model with M-CAP performed much better (>5% differences) for PCPG, TGCT and UVM cancer types, and lower for DLBC, CHOL, and SKCM than the models with Meta-LR, phastCons20way, and the constant scoring functions.

**Fig. 2.**
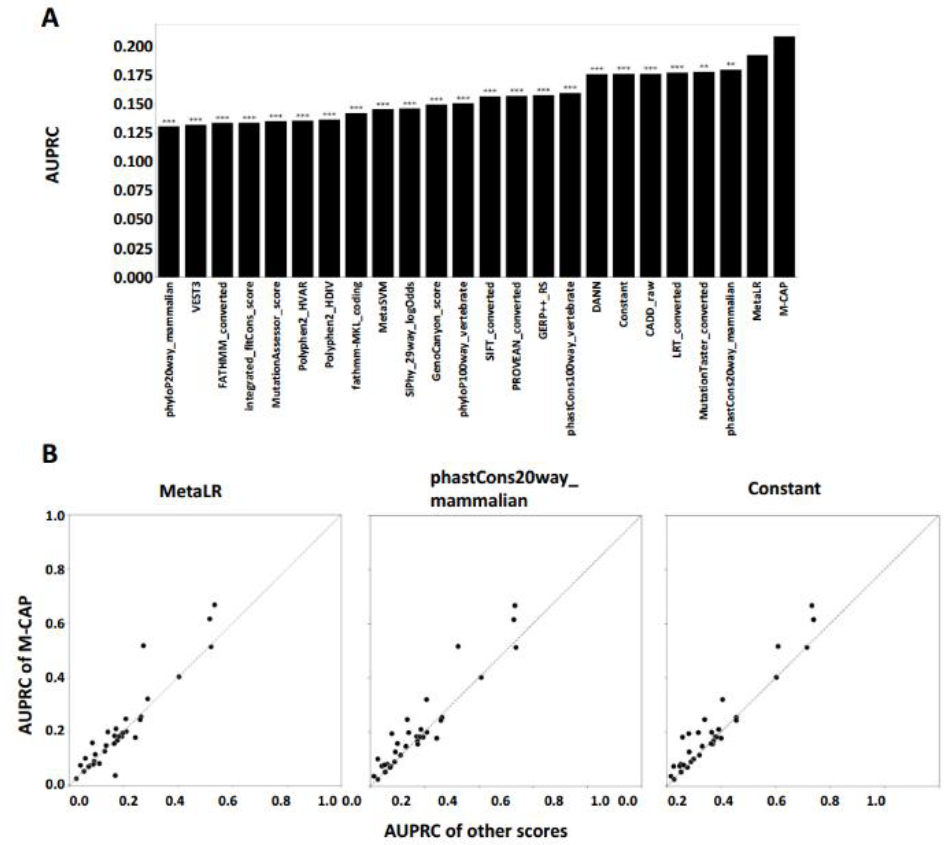
AUPRC comparison of different scoring systems in TCGA dataset. (A) M-CAP outperforms other 23 scoring systems in comparison of average AUPRC across 33 cancers. For each scoring system, the bar indicates the value of average AUPRC and the stars indicates its significance of mean difference compared with M-CAP. Mean differences were compared by paired t-test. ** and ***indicates P-value <0.01 and <0.001 respectively. (B) M-CAP generally performs better than other scoring functions in each cancer type. In scatter plots, each plot indicates AUPRCs against two scoring systems in the same cancer. Points above the diagonal line mean that AUPRCs of M-CAP are higher than that of others.

When counting the performing ranks of 24 scoring functions on 33 cancers (Table 1), the use of M-CAP ranked the 1^st^ in 9 cancers and within top 3 in 19 cancers. Although the use of MetaLR ranked the 1^st^ in 10 cancers, it ranked the top 3 for only 14. The use of the constant scoring function in DGAT-onco had the highest AUPRC for only two cancers. Thus, the M-CAP was used in our model if not specifically mentioned.

**Table 1.** The number of cancer types where each pathogenic score ranked the 1^st^, 2^nd^, and 3^nd^ in all scoring systems according to the AUPRC in the TCGA dataset.

| Score | 1<sup>st</sup> | 2<sup>nd</sup> | 3<sup>rd</sup> |
| --- | --- | --- | --- |
| MetaLR | 10 | 3 | 1 |
| M-CAP | 9 | 7 | 3 |
| phastCons20way_mammalian | 4 | 1 | 1 |
| SIFT_converted | 3 | 3 | 1 |
| LRT_converted | 2 | 1 | 1 |
| PROVEAN_converted | 2 | 3 | 2 |
| Constant | 2 | 3 | 4 |
| Others | 1 | 12 | 20 |

### 3.2 Comparisons with other methods in the TCGA and TS19 dataset

We evaluated the performances of 7 different methods (DGAT-onco, DiffMut, OncodriveCLUST, OncodriveFML, SomInaClust, WITER, and ITER) in both TCGA and the TS19 dataset. The TS19 dataset from other sources was selected as the independent test. As shown in Table 2, DGAT-onco has the highest average AUPRC in both TCGA and TS19 datasets (0.208 and 0.215 respectively), which are 17% and 5% higher than those of WITER, the 2^nd^ best method. As a frequency-based method without the integration of mutation functional impacts, DiffMut achieved the 3^rd^ best performance in the TCGA dataset but the worst performance in the TS19 dataset, likely because TS19 has relatively lower number of mutations (averagely 14674 mutations in TS19 compared to 48620 in TCGA of their corresponding 19 cancers). Paired t-tests indicated that DGAT-onco had signficantly higher AUPRC values than other methods except for WITER (Fig 3A) in the TCGA dataset, and better ones except for WITER and ITER in the TS19 dataset (Fig 3A). Fig S2 and S3 detailed the corresponding PRC (Precision-Recall curve) and AUPRC values.

**Table 2.**
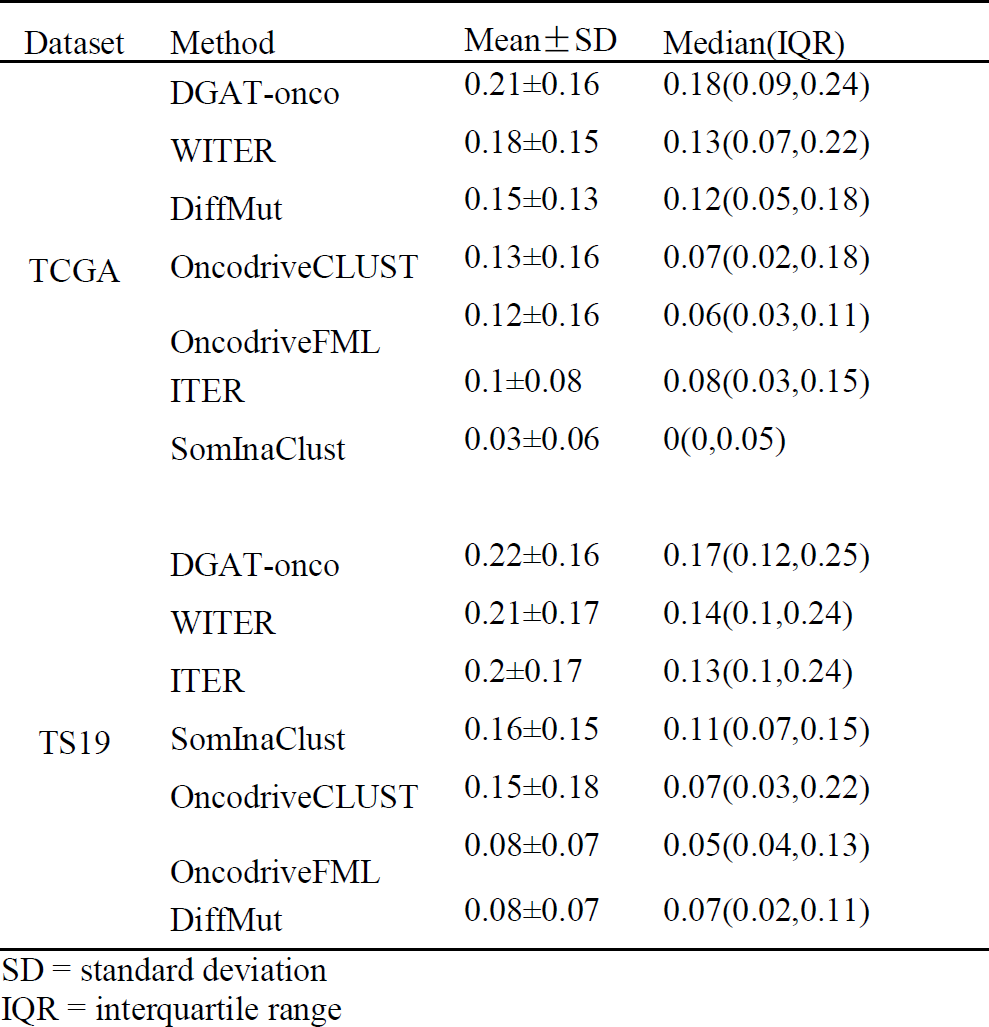
The comparison of average AUPRC values by DGAT-onco and other methods in the TCGA dataset.

| Dataset | Method | Mean ± SD | Median(IQR) |
| --- | --- | --- | --- |
| TCGA | DGAT-onco | 0.21±0.16 | 0.18(0.09,0.24) |
|  | WITER | 0.18±0.15 | 0.13(0.07,0.22) |
|  | DiffMut | 0.15±0.13 | 0.12(0.05,0.18) |
|  | OncodriveCLUST | 0.13±0.16 | 0.07(0.02,0.18) |
|  | OncodriveFML | 0.12±0.16 | 0.06(0.03,0.11) |
|  | ITER | 0.1±0.08 | 0.08(0.03,0.15) |
|  | SomInaClust | 0.03±0.06 | 0(0,0.05) |
| TS19 | DGAT-onco | 0.22±0.16 | 0.17(0.12,0.25) |
|  | WITER | 0.21±0.17 | 0.14(0.1,0.24) |
|  | ITER | 0.2±0.17 | 0.13(0.1,0.24) |
|  | SomInaClust | 0.16±0.15 | 0.11(0.07,0.15) |
|  | OncodriveCLUST | 0.15±0.18 | 0.07(0.03,0.22) |
|  | OncodriveFML | 0.08±0.07 | 0.05(0.04,0.13) |
|  | DiffMut | 0.08±0.07 | 0.07(0.02,0.11) |

SD = standard deviation IQR = interquartile range

**Fig. 3.**
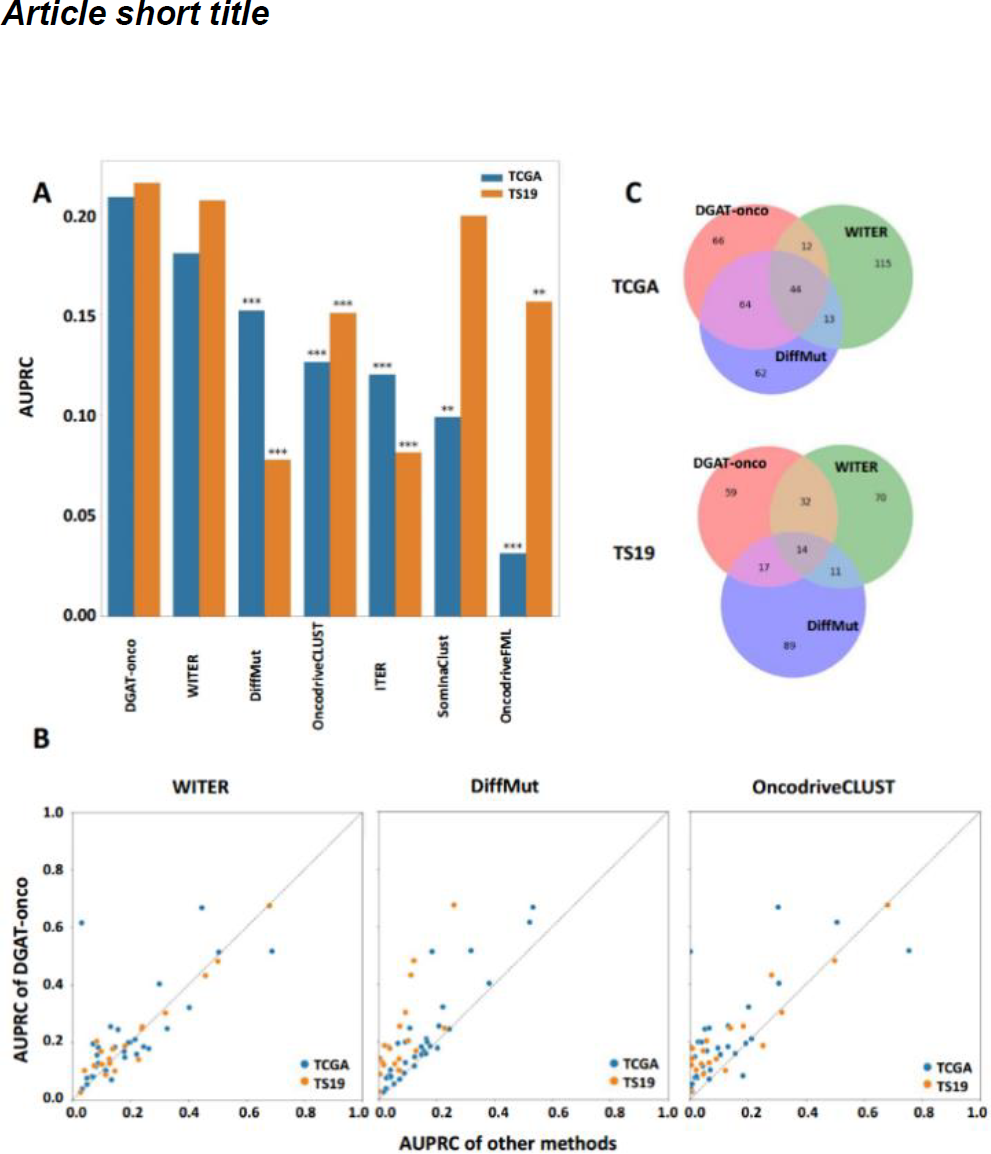
AUPRC comparison of difference methods. (A) DGAT-onco outperforms other 6 methods in comparison of average AUPRC. For each method, the bar indicates the value of average AUPRC and its significance of mean difference compared with DGAT-onco. Mean difference were compared by paired t-test. ** and ***indicates P-value <0.01 and <0.001, respectively. (B) DGAT-onco generally performs better in each cancer type. In scatter plots, each plot indicates AUPRCs against two methods in the same cancer. Points above the diagonal line mean that AUPRCs of DGAT-onco are higher than that of others. (C) DGAT-onco identidies different genes. For 3 methods, top 10 genes detected in each cancer are merged to a set. Venn plots show the relations of genes detected by 3 methods.

Pair-wise comparison between DGAT-onco and 3 representative methods (WITER, DiffMut, OncodriveCLUST) were detailed in scatter plots (Fig. 3B). In both datasets, DiffMut was overtaken by DGAT-onco in most cancers, while WITER and OncodriveCLUST have their own advantages and disadvantages in different cancers. As for OncodriveCLUST, it achieves good performance in THCA and UVM (AUPRC greater than 0.3) and performs badly (AUPRC less than 0.05) in several cancers (KIRC, LGG, MESO, PRAD, STAD). Although the average AUPRC of WITER is lower than that of DGAT-onco, points were surrounding near the diagonal line, and the classification superiority is not as clear as other methods.

We further evaluated the precision of different methods. The precision between top 1 to 50 genes of each cancer of TCGA and TS19 dataset were measured and details in Fig. S4 and Fig. S5. The average precisions across all cancer in these two dataset are shown in Fig. 4. Overall, in TCGA dataset, DGAT-onco has the highest precision, followed by WITER and DiffMut. In TS19 dataset, since WITER and ITER have the same precision as DGAT-onco in top 1 genes, they are overtaken as the threshold widens. The performance of DiffMut also drops sharply in the independent dataset. Besides,, when we merge top10 genes of each cancer detected by three methods to three set and found that, although the average precision and AUPRC between DGAT-onco and WITER were similar, they do identify a different set of potential oncogenes (Fig. 3C). Based on a complementary principle to identify oncogenes, DGAT-onco could select genes that may be missed by other methods.

**Fig. 4.**
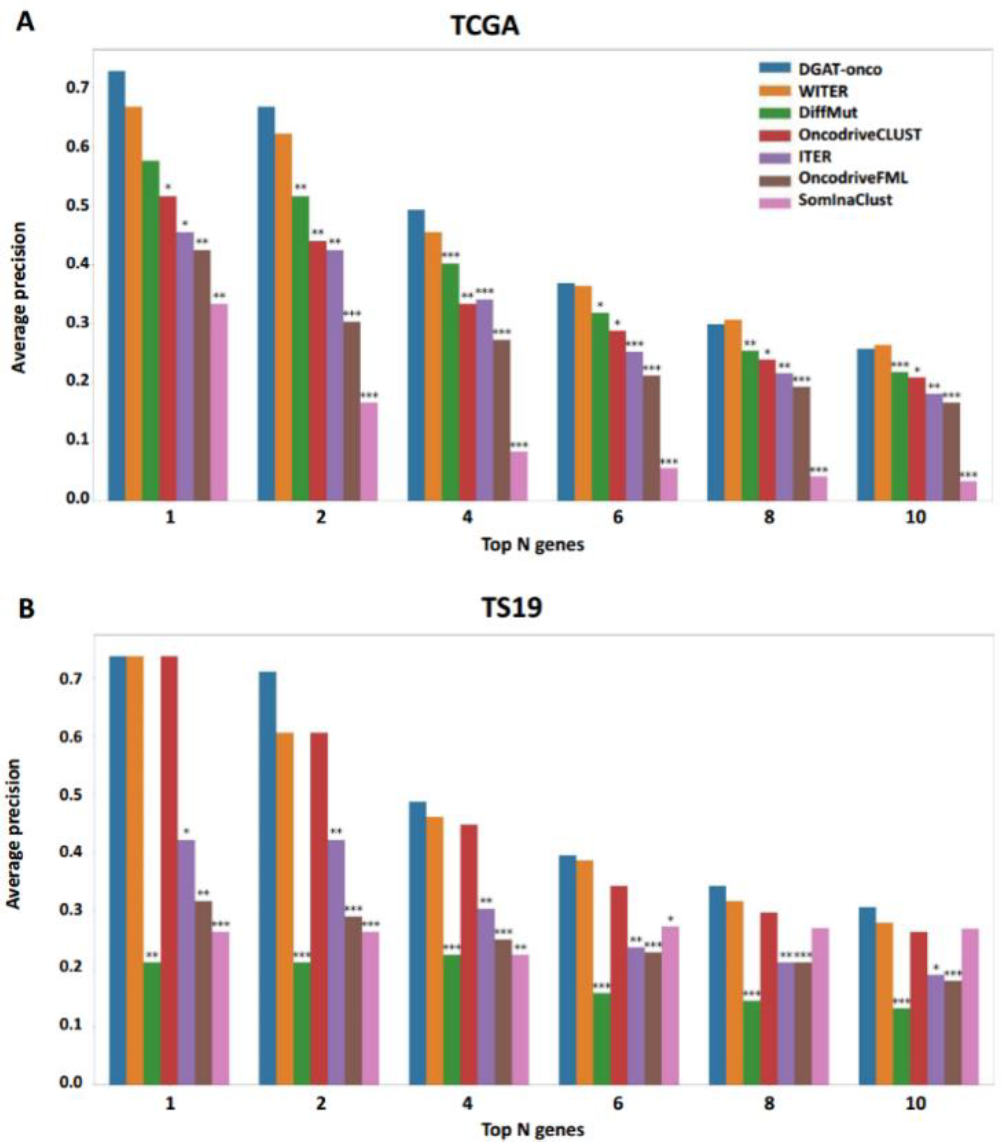
Precision comparison of different methods in TCGA and TS19 dataset. DGAT-onco generally outperformed other methods in the precision of top 10 genes in both dataset. For each method, the bar indicates the value of average precision and the stars indicates its significance of mean difference compared with DGAT-onco. Mean difference were compared by paired t-test. *, **, and ***indicates P-value <0.05, <0.01, and <0.001, respectively.

### 3.3 Functions of oncogenes predicted by DGAT-onco

DGAT-onco’s effectiveness provides a comprehensive landscape of oncogenes in multiple cancer types. Based on the analysis in TCGA dataset, among genes with top 50 uEMDs in each cancer, a total of 840 significant genes from 33 cancers were derived as potential oncogenes. The method detected 5 or more significant genes in 26 cancers, and out of which 22 cancers had more than 10 genes. Among these genes, 22.8% were verified as known oncogenes by CGC database. Details of significant genes and verified true oncogenes were shown in Supplementary3. The landscape of these genes was shown in a circos plot (Fig. 5A) (Gu, et al., 2014). In the plot, the length of the ring indicates the number of overlapped genes and the line represents their overlapping relationship. 30 out of 33 cancers have overlapped genes with other cancers. Ignoring cancer with equal to or less than 5 significant gene, COAD, BLCA and HNSC have the largest number of simultaneously significant genes with other cancers while COAD, CHOL and BRCA have the largest proportion of simultaneously significant genes (Fig. 5A). As expected, TP53 was the most common significant gene (in 22 cancers), followed by PIK3CA (17 cancers) and KRAS (12 cancers). Cancer with more significant genes was inclined to have more specific genes and implied high heterogeneity.

**Fig. 5.**
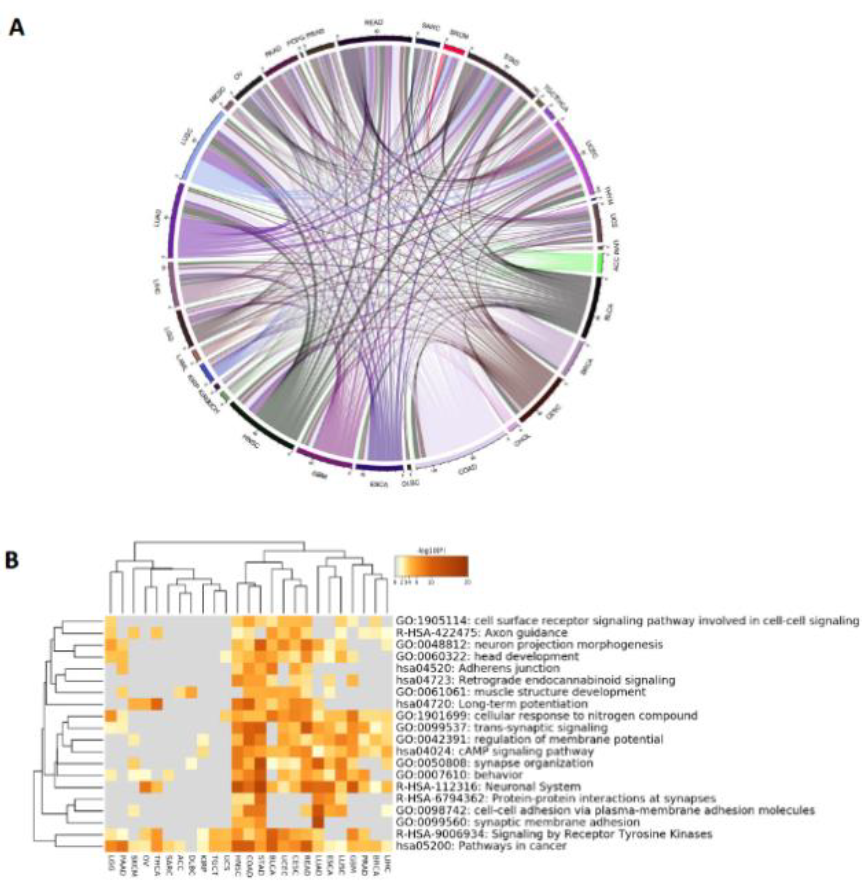
Oncogenes detected by DGAT-onco. (A) Circos plot of 840 significant genes in 33 cancers. The ring represents the number of overlapped significant genes with other cancers in a specific cancer type and the line represents the overlapping relationship between two cancers. (B) Significant genes were enriched in cancer-related pathways. Each row is a pathway or network and each column is a cancer set; the color in each cell indicates corresponding statistical significant (gray color indicates non-significant).

We further performed enrichment analysis to access the function of oncogenes predicted by DGAT-onco. After removing 3 cancers (CHOL, DLBC, UCS) with the lowest enrichment categories, 811 genes from 30 cancers were input for category search. The top enriched pathways of different cancer types were visualized as a heatmap (Fig. 5B). *Pathways in cancer* are the most common category among all cancer datasets, followed by *signaling by receptor tyrosine kinases* and *cellular response to nitrogen compound*. These functional categories are all cancer-related. The receptor tyrosine kinases are a major type of cell-surface receptors. Its signaling pathways have a various number of functions in cell proliferation, differentiation and survival. Some of receptor tyrosine kinases pathways have been described to be related with human cancers. As for nitrogen compound, since some of which have been prove to be carcinogens (Ward, 2009), the change of activity of cells it caused may be a signature of cancer development.

### 3.4 Case study: Predicting oncogenes in the ovarian cancer

We further selected the ovarian cancer (OV) in the TCGA dataset as a case to generate a set of predicted oncogenes. The detailed functions and roles of these genes in the cancer were obtained from the GeneCards database (https://www.genecards.org/), PubMed Gene database (https://www.ncbi.nlm.nih.gov/gene/), CGC database (https://cancer.sanger.ac.uk/census), and literature reviews. As shown in Table 3, among 16 significant genes, 5 genes were reported to be associated with cancers and 4 genes were reported to be associated with multiple cancers. In the CGC database, 4 genes were marked as cancer-related genes, and 3 were oncogenes related with multiple cancers.

**Table 3.**
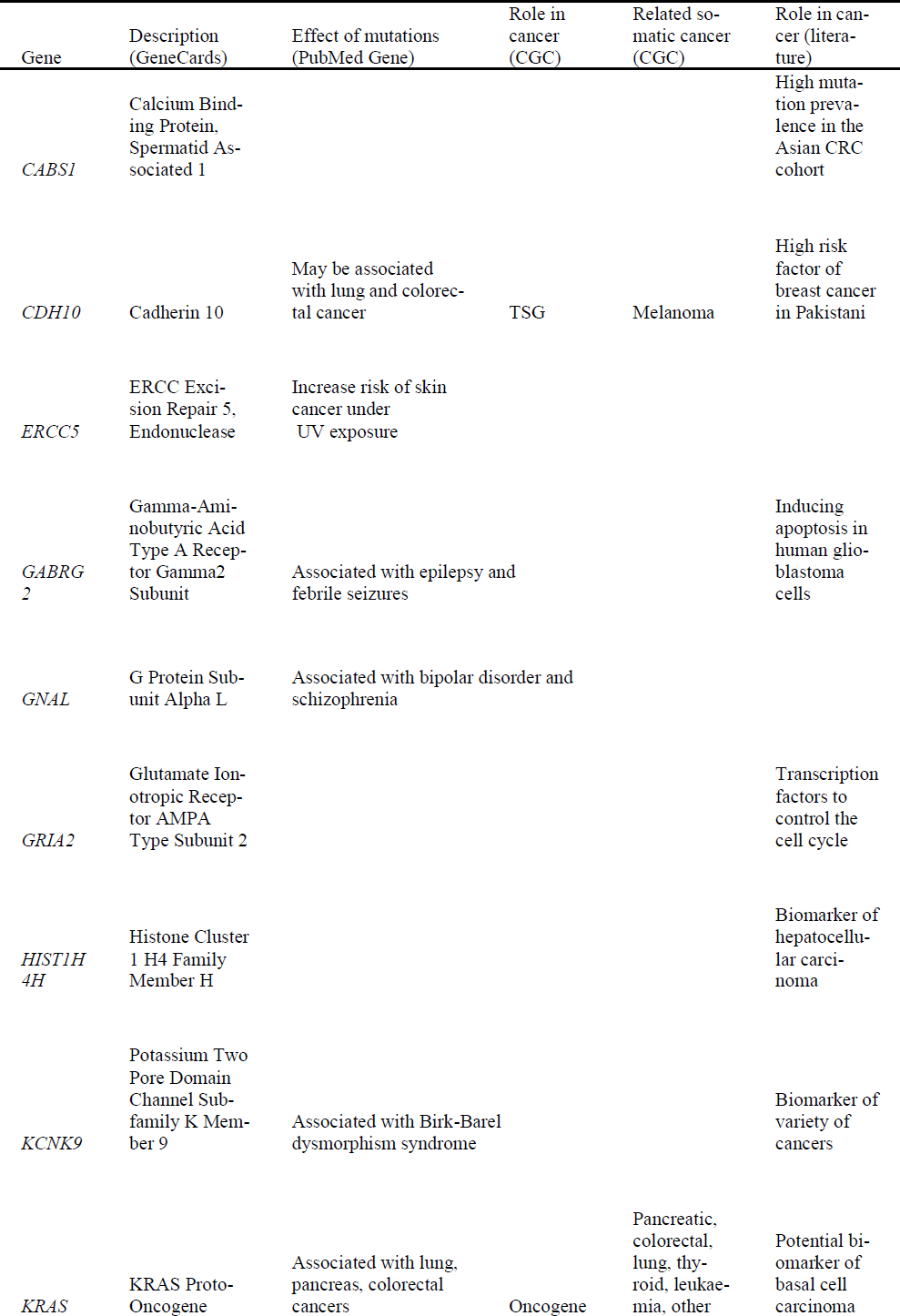
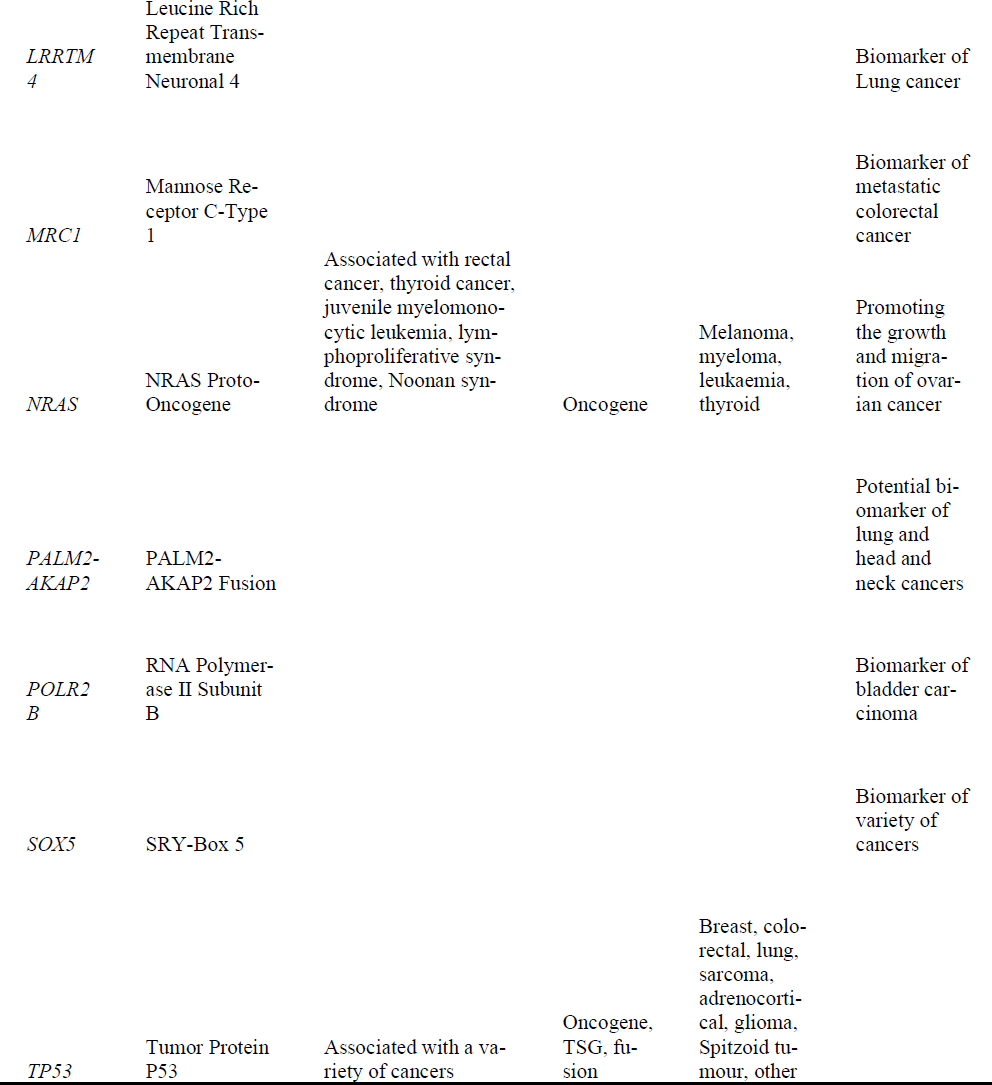
Description of significant genes in OV detected by DGAT-onco.

Literature reviews showed that 14 genes have were related with cancers (Ahn, et al., 2019; Chiou, et al., 2019; Ishiuchi, et al., 2002; Li, et al., 2019; Li, et al., 2016; Li, et al., 2017; Liu, et al., 2019; Malik, et al., 2019; Michiels, et al., 2007; Oakes, et al., 2017; Wang, et al., 2019; Yao, et al., 2018; Yu, et al., 2015). Among these genes, *KRAS, NRAS* and *TP53* are common oncogenes in cancer studies, while *CDH10, GABRG2, GRIA2, KCNK9, SOX5, ERCC5, HIST1H4H, LRRTM4* were recently verified as cancer-related gene by wet-lab experiments. The rest three genes (*MRC1, PALM2-AKAP2, POLR2B*) were discovered by other bioinformatics analysis study.

## 4 Discussion

The systematic genetic source from TCGA has initiated the wide practice of oncogene detection. Meanwhile, dbNSFP database including over 20 pathogenic systems for mutations makes it possible to generate pathogenic score profiles for various mutations. Moreover, the current method DiffMut was proved to have excellent performance compared with previous methods, but it does not efficiently use the pathogenic score systems. Pathogenic scores are not only important in distinguishing driver mutations from passenger ones, but also offers a reliable filtering criteria for oncogene detection. This motivated us to develop an improved method to integrate the useful pathogenic score features from available platforms in predicting oncogenes.

The classical methods in predicting oncogenes were based on classifying mutations types, like missense or nonsense, and might filter down some of them in a step. However, these methods ignored that even in missense mutations, their pathogenic effects can be quite different. Thus, it is necessary to employ the pathogenicity classifier to interpret the functional impacts of mutations. This study utilized the functional impacts of the mutations to detect the oncogenes and received an encouraging outcome. Although many previous methods have also realized the importance of mutation function impacts measurement, but most of them are limited in earlier pathogenic score like SIFT, PolyPhen-2, CADD (Kircher, et al., 2014). Our method reviewed the effectiveness of nearly all scoring systems in oncogenes detection systematically, including the latest systems had or had not been used by other methods. Besides, for background mutation modelling, we considered both the mutation number and their functional impacts in natural population, which offer a more accurate estimation than other methods. Although different pathogenicity score may focus on different purposes, 6 out of 23 pathogenicity score can contribute in oncogene prediction. In this study, the pathogenic score information was extracted from dbNSFP3.0 database, but any other source can be used for this purpose. In the future, as the development of better variant pathogenicity classifiers, the predictive performance of our method will keep increasing based on their knowledge.

Among 33 pathogenic scores, M-CAP shows the best performance in AUPRC measurement. M-CAP is a mutation scoring system developed on multiple pathogenic systems. As it was reported (Jagadeesh, et al., 2016), M-CAP can be used to interpret mutation linked to Mendelian diseases and has the ability to exceed other scores in pathogenic variants classification. It is well known that Mendelian disorders play an important role in genetic diseases caused by germline mutations like cystic fibrosis and Duchenne muscular dystrophy (Dietz, 2010). Although cancer is a genetic disease result from somatic mutations in the lifetime of a patient, genetic relationships between Mendelian disorders and cancer had been found and reported (Melamed, et al., 2015), which may explain the power of M-CAP in DGAT-onco.

To ensure that the classification superiority of DGAT-onco can be applied to genomics source other than TCGA, we selected an independent test dataset from other sources. The results clearly show that DGAT-onco outperforms alternative methods. Two frequency-based methods, DiffMut and ITER, are all overtaken by their improved versions (DGAT-onco and WITER) that integrated function impacts of mutations. Since the number of samples and mutations in the independent test dataset are much less than that of TCGA, the performance of DiffMut, drops significantly in the independent test dataset. At the same time, WITER and ITER, methods follows step-wise procedure to detect oncogenes from a mutation model, have advantages in the independent test dataset that has a small sample size.

Besides, the enrichment results also showed that the genes identified by DGAT-onco are highly related with cancer. Apart from the most common category (pathways in cancer), pathways related with signaling, cell differentiation and impacts of compound have been identified. Since cancer is disease initiated by out of control, potential risk factors such as ignoring signals to differentiate, damage in important signaling pathways and accumulation of contaminants may play a role in cancer development.

Currently, DGAT-onco does not consider the mutations in introns such as 3-UTR and 5-UTR because no M-CAP score annotations were conducted in these regions. Nevertheless, due to the potential contributions of non-coding mutations on cancer (Fredriksson, et al., 2014; Puente, et al., 2015), the development of new metric aimed at non-coding mutations may improve the performance of DGAT-onco. Another option to solve this issue may be summarizing the information of this region by frequency-based methods without pathogenicity score, but the detail of combination between the non-coding region and coding region need to be considered carefully. Additionally, since cancer is a disease that may be contributed by a set of genes (Creixell, et al., 2015), it is critical to interpreting its mechanisms by biological pathways and protein-protein interactions (Kar, et al., 2009). However, DGAT-onco is a single gene comparison method neglecting the effect or situation of other genes, which the features are not fully used in such a framework. Using a network-based method to integrate the known interaction network or conducting genes cluster member search to making the results more interpretability may improve the performance of DGAT-onco.

## Supporting information

all supplementray

## Acknowledgements

Our method is based upon data generated by the TCGA Research Network (https://www.cancer.gov/tcga) and cBioPortal (https://www.cbioportal.org/). We thank DiffMut team for their enlightening work.

