## Supplementary material for "DGAT-onco: A powerful method to detect oncogenes by integrating differential mutational analysis and functional impacts of somatic mutations": all supplementray: Supplementary1.docx

**Notes**

**The usage of the alternative driver-gene estimation methods**

Our method was compared to 6 popular methods, including DiffMut, OncodriveCLUST, OncodriveFML, SomInaClust, WITER and ITER. The DiffMut, OncodriveCLUST, and SomInaClust were run by their R packages. The WITER and ITER were run on their JAVA tools. Since OncodriveFML only accepted MAF of the hg19 version, our MAF of the hg 38 version was converted to hg 19 version through the CrossMap (Zhao, et al., 2013) and then submitted to the webserver (<http://bbglab.irbbarcelona.org/oncodrivefml/home>). For all methods, the default or suggested parameters were used excepted we meet a program error caused by limited mutation number.

For 3 cancers in OncodriveCLUST (2 in the TCGA dataset, 1 in the TS19 dataset) with a limited number of mutation, the program went wrong when we used the default parameters, so we only changed the “minMut” arugement to 1(default 5) for these 3 datasets to avoid program errors.

For 12 cancers in WITER (3 in the TCGA dataset, 4 in the TS19 dataset ) and ITER (1 in the TCGA dataset, 4 in the TS19 dataset ) with a limited number of mutation, we use WITER-Ref (WITER with a BRCA reference for background gene mutation) to generated resultes.

**The measurement of AUPRC**

The statistics of DGAT-onco, DiffMut, OncodriveCLUST, OncodriveFML, SomInaClust, WITER and ITER were uEMD, uEMD, clustering score, qDG, OG score, q value and q value respectively. The gold standard of an oncogene is that it has been marked as an oncogene according to the somatic mutations in the CGC database.

With respect to the plotting of PRC, we first used each sorted statistics as a threshold to measure its precision and recall. Then, we connected every two points from left to right by a smooth line. To do this smooth line, we evenly split the difference of recalls between two points and calculated the corresponding precisions, before connecting these points. If a curve had not point at y-axis, the leftmost point was copied and moved to the y-axis.

With respect to the plotting of measurement of AUPRC, we calculated the area between two points one by one from left to right before summing up. If twos points have the same precision, we calculated area according to rectangle area formula. If twos points have different precision, we calculated the area by: $\sum_{i=1}^{n} \frac{{TP}_{i}}{{TP}_{i}+{FP}_{i}}*\Delta\frac{{TP}_{i}}{Pos}$ , where ${TP}_{i}= {TP}_{l}+{(TP}_{r}-{TP}_{l})*\frac{i}{n}$ , ${FP}_{i}= {FP}_{l}+({FP}_{r}-{FP}_{l})*\frac{i}{n}$, n is a number approaching infinity, *TP_l_* and *TP_r_* are the number of true positive at left and right point, *FP_l_* and *FP_r_* are the number of false positive at left and right point, *Pos* is the number of positive samples (known oncogenes). After changing the expression to $\int_{0}^{1} \frac{{TP}_{l}+({TP}_{l}-{TP}_{r})x}{{TP}_{l}+{FP}_{l}+({TP}_{r}+{FP}_{r}-{TP}_{r}-{FP}_{r})x}*\frac{{(TP}_{r}-{TP}_{l})dx}{Pos}$ , we can obtain the area between these two points.

**Table S1**. Summary of datasets used in the study

| dataset | cancer | samples (n) | genes (n) | mutations (n) | nonsynonymous  mutations (n) | source |
| --- | --- | --- | --- | --- | --- | --- |
| TCGA | ACC | 92 | 5441 | 7704 | 4337 | TCGA |
|  | BLCA | 408 | 17283 | 86284 | 50675 | TCGA |
|  | BRCA | 970 | 16320 | 60823 | 35197 | TCGA |
|  | CESC | 288 | 17646 | 72389 | 35779 | TCGA |
|  | CHOL | 49 | 2675 | 3062 | 1468 | TCGA |
|  | COAD | 399 | 18916 | 156276 | 90484 | TCGA |
|  | DLBC | 37 | 3267 | 4294 | 2428 | TCGA |
|  | ESCA | 182 | 11905 | 26864 | 13668 | TCGA |
|  | GBM | 386 | 13667 | 38573 | 23298 | TCGA |
|  | HNSC | 505 | 15680 | 60475 | 36314 | TCGA |
|  | KICH | 66 | 1569 | 1711 | 987 | TCGA |
|  | KIRC | 335 | 8122 | 13250 | 8131 | TCGA |
|  | KIRP | 280 | 9308 | 16194 | 9566 | TCGA |
|  | LAML | 94 | 1148 | 1280 | 772 | TCGA |
|  | LGG | 499 | 10655 | 22022 | 13711 | TCGA |
|  | LIHC | 362 | 13942 | 37970 | 21787 | TCGA |
|  | LUAD | 554 | 17194 | 114390 | 71097 | TCGA |
|  | LUSC | 486 | 17470 | 113724 | 69500 | TCGA |
|  | MESO | 78 | 2067 | 2351 | 1318 | TCGA |
|  | OV | 432 | 11414 | 24749 | 15256 | TCGA |
|  | PAAD | 163 | 7763 | 12848 | 8102 | TCGA |
|  | PCPG | 174 | 1127 | 1248 | 787 | TCGA |
|  | PRAD | 462 | 9431 | 17396 | 10680 | TCGA |
|  | READ | 136 | 14232 | 44600 | 26046 | TCGA |
|  | SARC | 236 | 9660 | 17959 | 8782 | TCGA |
|  | SKCM | 465 | 18869 | 300132 | 163453 | TCGA |
|  | STAD | 427 | 17788 | 112306 | 68570 | TCGA |
|  | TGCT | 139 | 1361 | 1513 | 908 | TCGA |
|  | THCA | 477 | 2863 | 3775 | 2442 | TCGA |
|  | THYM | 95 | 1754 | 1996 | 1096 | TCGA |
|  | UCEC | 528 | 20933 | 637224 | 331479 | TCGA |
|  | UCS | 57 | 5516 | 7769 | 4834 | TCGA |
|  | UVM | 80 | 1069 | 1258 | 839 | TCGA |
| TS19 | BLCA | 109 | 235 | 1177 | 819 | Kim et al. Eur Urol 2015 |
|  | BRCA | 2369 | 173 | 17272 | 10165 | Pereira et al. Nat Commun 2016 |
|  | CHOL | 182 | 237 | 767 | 467 | Lowery et al. Clin Cancer Res 2018 |
|  | COAD | 106 | 16240 | 73833 | 41998 | Vasaikar et al. Cell 2019 |
|  | DLBC | 972 | 150 | 7105 | 5751 | Reddy et al. Cell 2017 |
|  | ESCA | 146 | 12246 | 31023 | 17036 | Dulak et al. Nat Genet 2013 |
|  | HNSC | 74 | 6055 | 9603 | 6262 | Stransky et al. Science 2011 |
|  | KIRC | 81 | 354 | 532 | 349 | Sato et al. Nat Genet 2013 |
|  | LAML | 608 | 3335 | 9910 | 7664 | Tyner et al. Nature 2018 |
|  | LGG | 61 | 8447 | 15155 | 8886 | Johnson et al. Science 2014 |
|  | LIHC | 243 | 11834 | 28170 | 16941 | Schulze et al. Nat Genet 2013 |
|  | LUAD | 183 | 14770 | 65767 | 42982 | Imielinksi et al. Cell 2012 |
|  | PAAD | 383 | 10498 | 23536 | 13372 | Bailey et al. Nature 2016 |
|  | PCPG | 1012 | 15827 | 95337 | 51717 | Armenia et al. Nat Genet 2018 |
|  | SKCM | 91 | 9104 | 26169 | 23601 | Krauthammer et al. Nat Genet 2012 |
|  | STAD | 100 | 8749 | 17901 | 13042 | Wang et al. Nat Genet 2014 |
|  | UCEC | 195 | 362 | 2545 | 1775 | Soumerai et al. Clin Cancer Res. 2018 |
|  | UCS | 22 | 9728 | 19148 | 15735 | Jones et al. Nat Commun 2014 |
|  | UVM | 25 | 255 | 299 | 244 | Johansson et al. Oncotarget 2016 |
| 1000G |  | 2504 | 17891 | 84739838 | 569128 | 1000G |

**Table S2.** Summary of pathogenicity scoring systems used in the study

| Score | Training source | Prediction method |
| --- | --- | --- |
| CADD_raw | Observed and simulated variants | Support vector machine |
| DANN | Observed and simulated variants | Deep neural network |
| FATHMM_converted | SNVs from HGMD and UniProt | Hidden Markov models |
| fathmm-MKL_coding | SNVs from HGMD and 1000G | Multiple kernel learning |
| GenoCanyon | Computational and experimental annotations | Unsupervised learning |
| GERP++_RS | Genomes of mammals | Maximum likelihood estimation |
| integrated_fitCons | Genomes of unrelated human | Other |
| LRT_converted | Coding sequences of vertebrate species | Likelihood ratio test |
| M-CAP | Multiple pathogenicity scores | Gradient boosting trees |
| MetaLR | SNVs from UnProt | Logistic regression |
| MetaSVM | SNVs from UnProt | Support vector machine |
| MutationAssessor | SNVs from COSMIC | Other |
| MutationTaster_converted | SNVs from 1000G and HGMD | Naive Bayes classifier |
| phastCons100way_vertebrate | Genomes of vertebrates | Hidden Markov model |
| phastCons20way_mammalian | Genomes of mammals | Hidden Markov model |
| phyloP100way_vertebrate | Genomes of vertebrates | Hidden Markov model |
| phyloP20way_mammalian | Genomes of mammals | Hidden Markov model |
| Polyphen2_HDIV | SNVs from close mammaliam | Naive Bayes classifier |
| Polyphen2_HVAR | Disease-associated SNVs and common SNVs | Naive Bayes classifier |
| PROVEAN_converted | SNVs from UniProt and HUMSAVAR | Other |
| SIFT_converted | Deleterious and tolerant nsSNVs of E. coli LacI | Position-specific scoring matrix |
| SiPhy_29way_logOdds | Genomes of mammals | Other |
| VEST3 | SNVs from HGMD and the exome sequencing | Random forest |

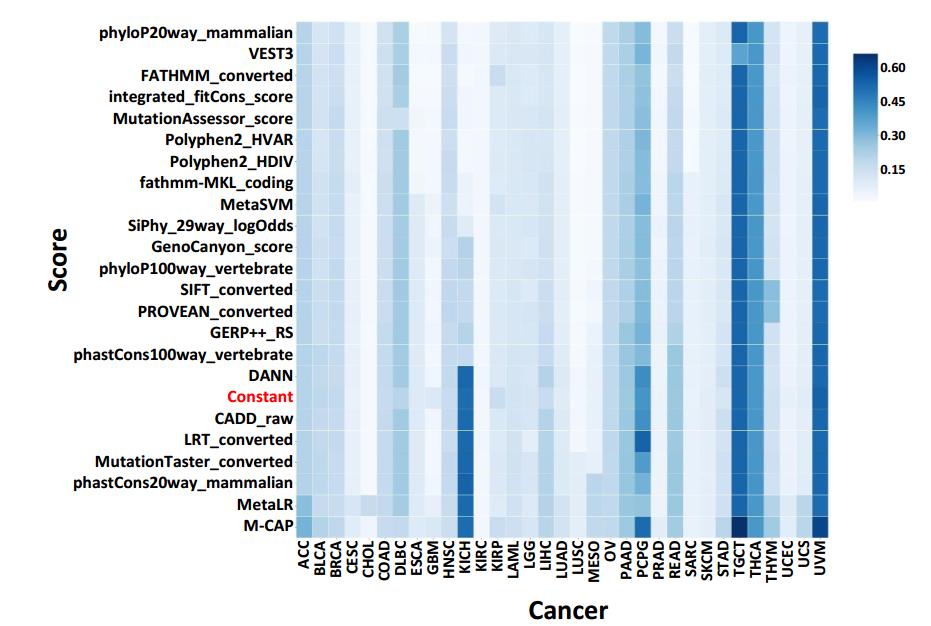

**Fig. S1** DGAT-onco models comparison in TCGA dataset. The colour depth indicates the magnitude of AUPRC. The rows are scores of models and they are sorted by the average AUPRC. The columns indicate different cancers.

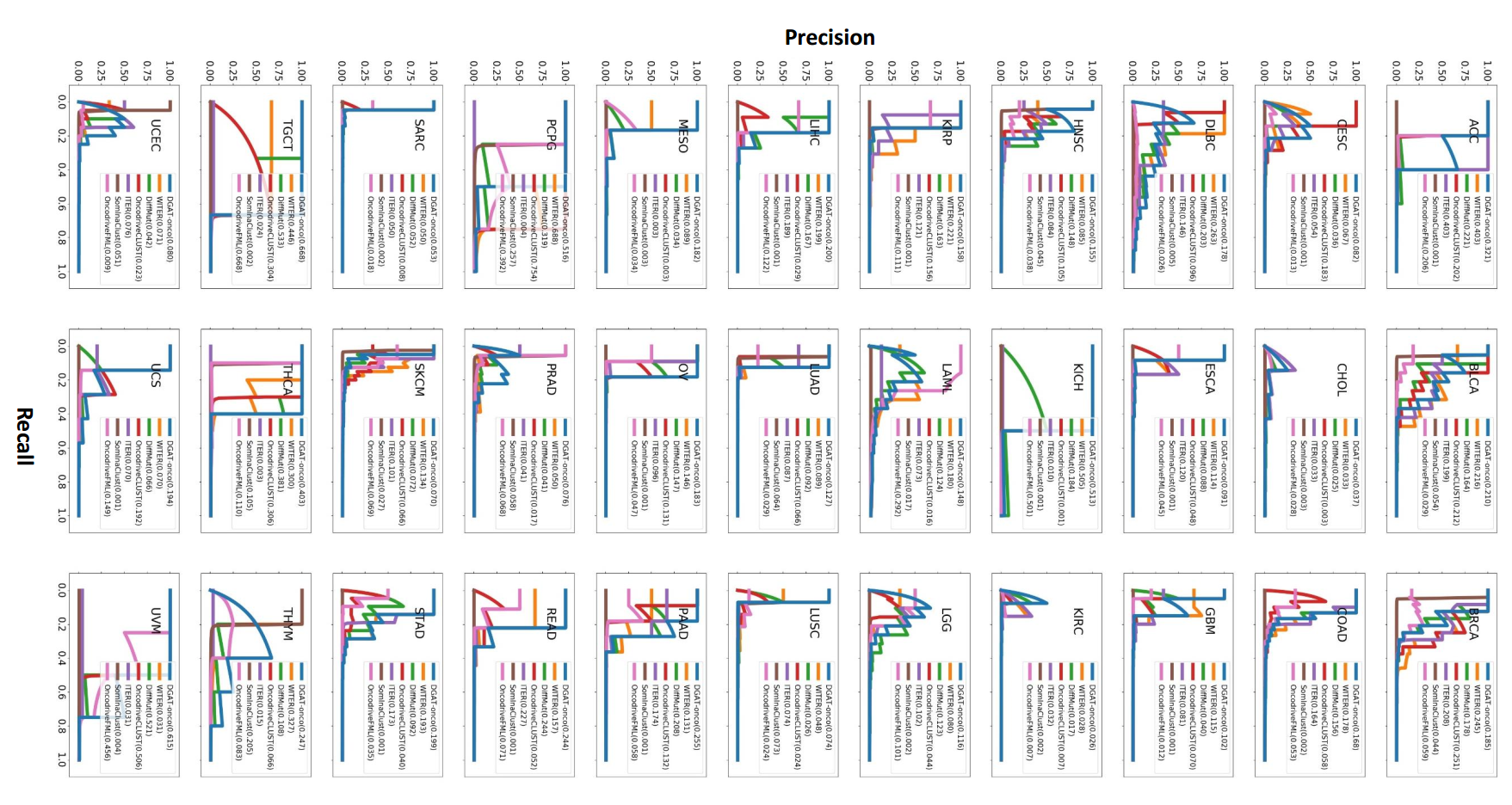

**Fig. S2.** PRC of all methods in TCGA dataset. If a method have no output for a dataset, the PRC is a horizontal line with the height equal to n (oncogenes)/ n (total genes).

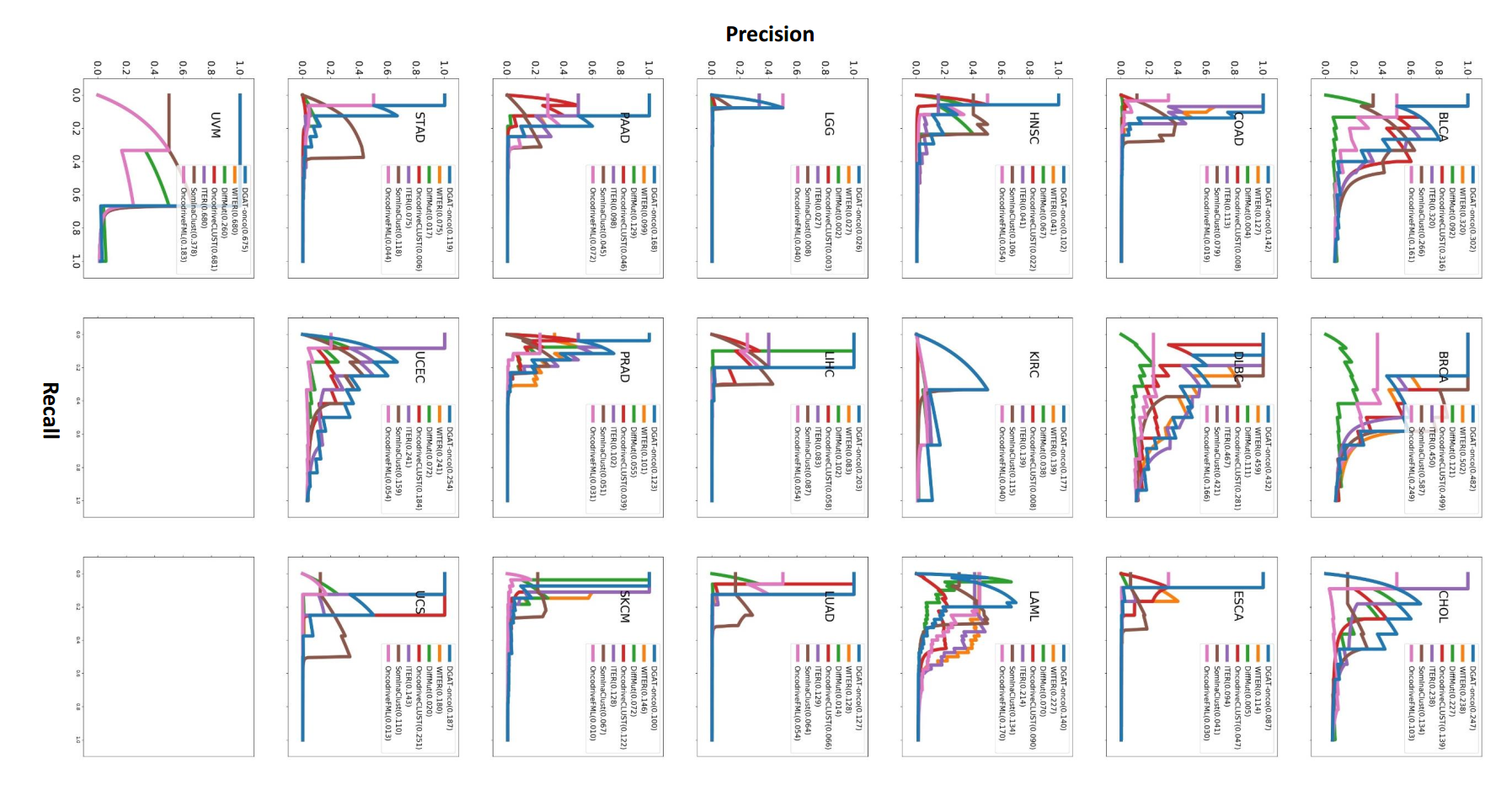

**Fig. S3.** PRC of all methods in the TS19 dataset. If a method have no output for a dataset, the PRC is a horizontal line with the height equal to n (oncogenes)/ n (total genes).

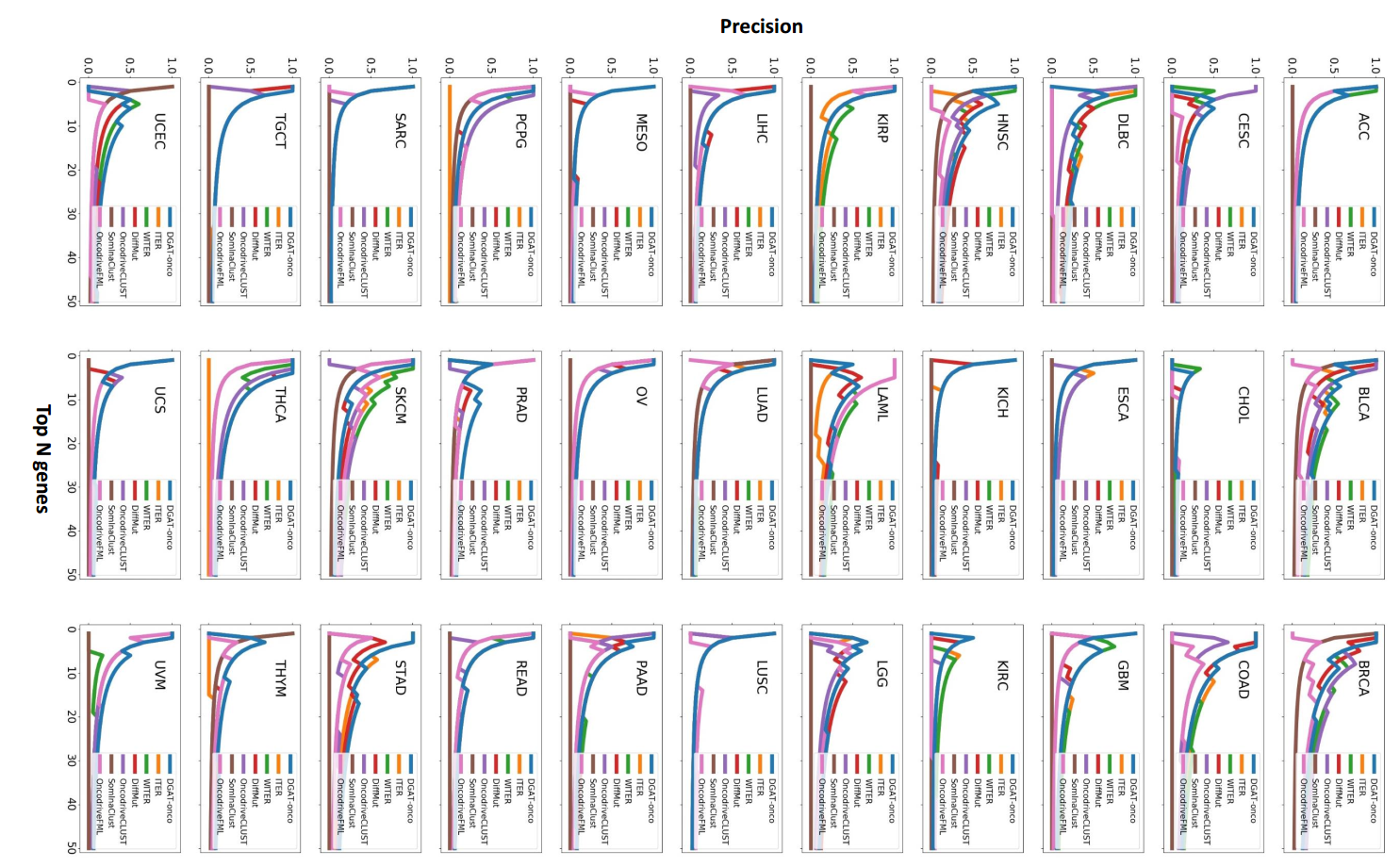

**Fig. S4.** Precision of top 50 genes of all methods in TCGA dataset. If a method have no output for a dataset, the curve is a horizontal line with the height equal to 0.

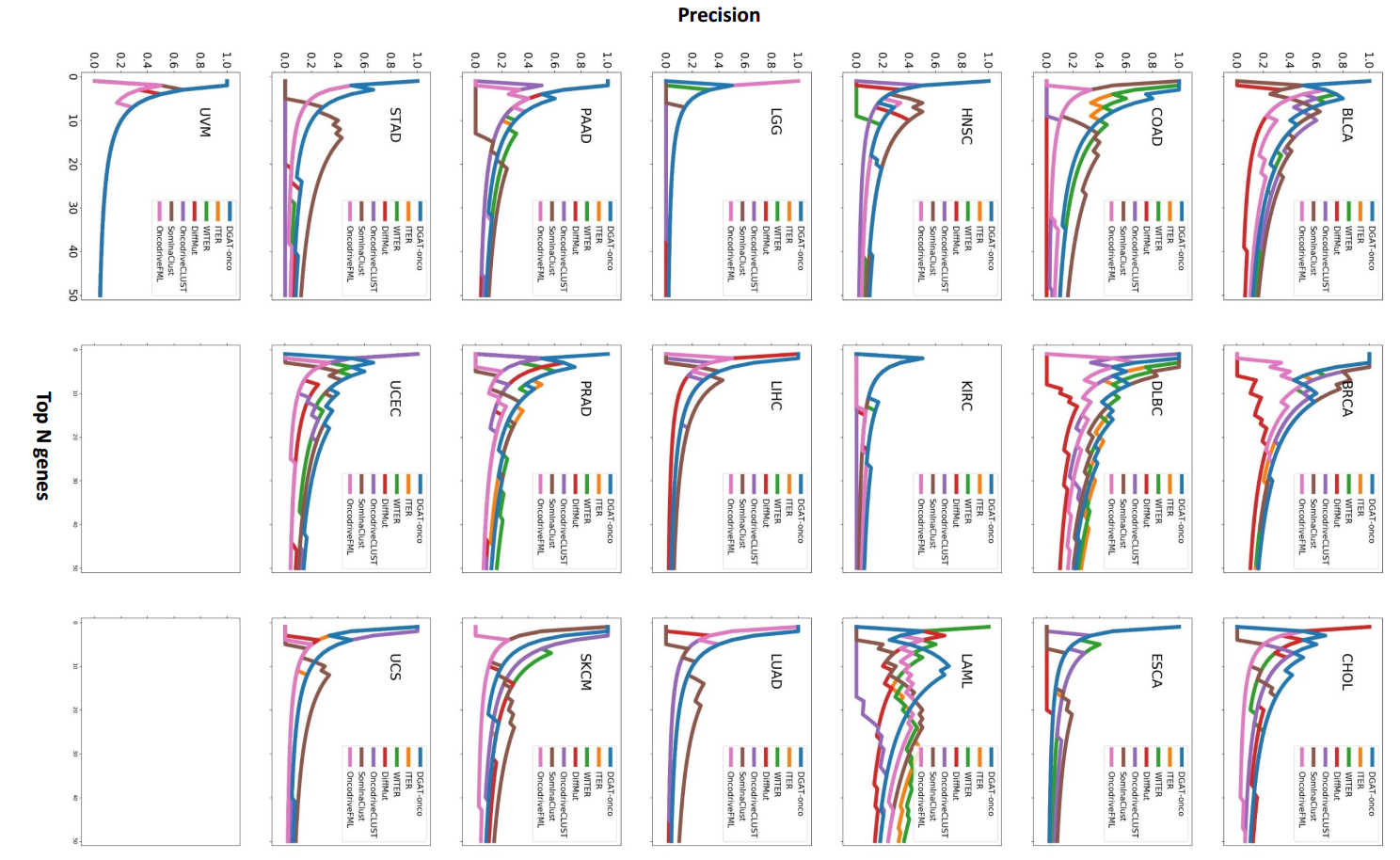

**Fig. S5.** Precision of top 50 genes of all methods in the TS19 dataset. If a method have no output for a dataset, the curve is a horizontal line with the height equal to 0.
